# The Effects of Practice-based Training on Graduate Teaching Assistants’ Classroom Practices

**DOI:** 10.1101/115295

**Authors:** Erin A. Becker, Erin J. Easlon, Sarah C. Potter, Alberto Guzman-Alvarez, Jensen M. Spear, Marc T. Facciotti, Michele M. Igo, Mitchell Singer, Christopher Pagliarulo

## Abstract

Evidence-based teaching is a highly complex skill, requiring repeated cycles of deliberate practice and feedback to master. Despite existing well characterized frameworks for practice-based training in K-12 teacher education, the major principles of these frameworks have not yet been transferred to instructor development in higher educational contexts, including training of graduate teaching assistants (GTAs). We sought to determine whether a practice-based training program could help GTAs learn and use evidence-based teaching methods in their classrooms. We implemented a weekly training program for introductory biology GTAs, which included structured drills of techniques selected to enhance student practice, logic-development, and accountability and reduce apprehension. GTAs received regular performance feedback based on classroom observations. To quantify use of target techniques and levels of student participation, we collected and coded 160 hours of video footage. We found that, although GTAs adopted and utilized many of the target techniques with high frequency, techniques which enforced student participation were not stably adopted and their use was unresponsive to formal feedback. We also found that techniques discussed in training, but not practiced, were not used at quantifiable frequencies, further supporting the importance of practice-based training for influencing instructional practices.

## Introduction

Introductory STEM courses at research-intensive universities tend to employ didactic, lecture-based instruction (Lund et al. 2015). This type of learning environment, on its own, offers few opportunities to engage in the kind of iterative practice and feedback necessary for students to test and improve their current state of knowledge. To provide opportunities for structured practice, many of these courses include an associated discussion or laboratory section, often taught by graduate teaching assistants (GTAs). Analogously, existing pedagogy training for GTAs is often structured as didactic workshops, peer discussions, and/or literature readings (Prieto and Scheel 2008), and often does not facilitate the type of iterative practice and feedback needed to learn and master such skills.

Here we focus on the development and implementation of a practice-based teacher training program for GTAs. Practice-based teaching frameworks are widely-used in K-12 teacher training (Zeichner 2012) with evidence showing that such “hands-on” work is an important element in the success of professional development in impacting teachers’ classroom practices (Garet et al. 2001). Such training engages novice teachers in deliberate practice (Ericsson, Krampe, and Tesch-Römer 1993) of well-specified instructional activities, and incorporates targeted feedback and coaching (Lampert et al. 2010). Training focuses on a core set of evidence-based high-impact teaching practices that occur with high frequency, can be used across many different environments, and which instructors can begin to master early in their teaching careers (Grossman, Hammerness, and Mcdonald 2009). Complex teaching practices are first broken down into learnable component skills. Novices are then provided with repeated and progressive practice of these skills to develop automaticity (Darling-Hammond and Bransford 2007). Focus on a limited number of component skills allows novices to build efficacy with demanding practices in small steps; this occurs through repeated cycles of practice in increasingly complex situations (Lampert et al. 2013). During practice, component skills are first modeled by trainers and then drilled in a simulated, simplified situation with peers. Finally, component skills are implemented in an authentic classroom environment, under close observation, with coaching.

Each of the instructional practices included in our GTA training program were selected for alignment with research-based dimensions of active learning, namely enhancing student accountability, practice, logic-development, and apprehension reduction (Eddy, Converse, and Wenderoth 2015). One mechanism which has been proposed to explain improvements in student learning outcomes observed in active learning environments, is that increased student participation in the learning process requires students to take responsibility for their own knowledge level (Eddy, Converse, and Wenderoth 2015). When participation is enforced, students should be more likely to hold themselves and each other accountable, leading to an increase in focused time on task. Based on this proposed mechanism, GTAs were instructed to call on every student during each class period. This constituted a “task-standard” for performance.

Weekly drills constituted the “task repetition” element of deliberate practice. In addition, we utilized a comprehensive feedback program, consistent with current recommendations for best practices (Gormally, Evans, and Brickman 2014). This program included timely and repeated feedback sessions (Fedor and Buckley 1987; O’Reilly and Renzaglia 1994; Rezler and Anderson 1971) focused on an explicit task-standard for performance (Hattie and Timperley 2007), and provided measurable and specific guidelines for improvement (Englert and Sugai 1983; Liden and Mitchell 1985; O’Reilly and Renzaglia 1994). Together, the task repetition and feedback elements of our training program fulfill the demonstrated requirements for effective deliberate practice (Ericsson, Krampe, and Tesch-Römer 1993) and closely align with practice-based training guidelines developed for K-12 teaching training.

Our study addressed two primary research questions. First, how did GTAs in a practice-based training program implement evidence-based instructional practices and was this implementation successful at eliciting high levels of student participation? Secondly, how did GTA practice change throughout the training program and were these changes associated with whether they were given feedback on their use of specific techniques? We also addressed a secondary research question - which, if any, of the targeted techniques were associated with changes in student learning outcomes?

Traditionally, the impact of GTA training programs have been assessed using indirect measures, such as self-efficacy surveys (Hardré 2003; Komarraju 2008; Young and Bippus 2008), student evaluations (Davis and Kring 2001; Marbach-Ad et al. 2012; Pentecost et al. 2012; Schussler et al. 2015) and surveys of perceived student learning (Hardré 2003). Use of such indirect measures can be valuable. However, by themselves, they do not provide detailed information about the actual use of instructional practices, and therefore cannot address the question of how training efforts translate into concrete changes in teaching behavior. Thus, it is crucial to measure GTA classroom practices directly, both to enable evaluation of the effectiveness of the training program, and to provide accurate feedback to trainees.

To assess the success of our training program, we evaluated three outcome variables, which align with three of Kirkpatrick’s four levels of evaluation (Kirkpatrick 1994). This framework has previously been used to evaluate professional development in higher education contexts (Steinert et al. 2006; Wyse, Long, and Ebert-May 2014). Our outcome variables were 1) GTA reaction to training program, 2) demonstration of GTAs’ ability to apply target instructional practices in the classroom, and 3) impact of training on student learning. These three outcome variables correspond to the three types of outcome variables in Reeve’s framework for graduate teaching assistant professional development evaluation and research: 1) GTA cognition, 2) GTA teaching and 3) undergraduate student outcomes (Reeves et al. 2016).

Our study is unique in implementing a GTA training program strongly aligned with practice-based frameworks found in K-12 teaching training, and in directly assessing both the impact of training on GTA classroom practices longitudinally throughout the training process and associations between GTA instructional practices and student learning.

## Methods

### Course structure

This study was conducted at a four-year, residential R1 research university in the Western United States. All 15 GTAs involved in the study led discussion sections for the same course, taught by the same lecture instructor in Fall 2014. Involvement in the training program was mandatory. The course was a large-enrollment (~1,000 students/quarter) first course in a three-quarter introductory general biology series. Course content focused on molecular and cellular biology, including: biochemistry, energetics, metabolism, cellular structure, information flow and regulation. The course met four times per week (three one-hour lecture periods and one two-hour GTA-lead discussion period) for 10 weeks. Student attendance at discussion was mandatory, with discussion scores comprising 20% of the final course grade. Content covered in each weekly discussion was standardized across all 45 discussion sections. A summary of topics covered each week is shown in **Supplementary Table 1**.

Prior to attending each weekly discussion, students completed graded pre-discussion quizzes (preparatory assignments) through Carnegie Mellon University’s Open Learning Initiative (OLI) Introduction to Biology online course (http://oli.cmu.edu/), modified to fit specific course content. These quizzes were accompanied by substantial reading material and ungraded practice problems that included immediate, automated feedback. Student preparatory assignment responses provided feedback to GTAs about their students’ level of understanding and were used by each GTA to design individualized 4560-minute question and answer warm-up sessions held at the beginning of each discussion. The remainder of class time (40-55 min) was spent on Process Oriented Guided Inquiry Learning (POGIL)-like problem sets focusing on case studies and guided inquiry activities (Farrell, Moog, and Spencer 1999) which were graded and served as weekly review assignments. Students worked in groups of four on problem sets and during group-based warm-up questions. During warm-up question and answer session, GTAs were free to use any instructional method they felt would best elicit student participation, however they were strongly encouraged to use the engagement techniques covered in training (detailed below). During the remainder of class time, GTAs were directed to move throughout the classroom and interact with student groups to gauge their level of understanding and to supply individualized feedback to students (“Circulate and Check for Understanding” – detailed below).

### GTA demographics

Information on GTA demographics and previous teaching experience was compiled from GTA self-reported data and university records (**Table 1**). Forty percent of GTAs had previously served as GTAs for the study course and sixty-six percent had acted as GTAs for some course prior to the study term. One third of GTAs had other (non-GTA) teaching experience prior to the study term, either at the university level or in other educational settings.

**Table 1:** GTA demographics. All teaching experience is prior to and not including study term. Mean and standard deviation for length of teaching experience do not include respondents who reported no experience in that category. N = 15 GTAs.

|  | value | % responding |
| --- | --- | --- |
| % female | 33% | 100 |
| % domestic | 87% | 100 |
| % in PhD program | 73% | 100 |
| Year in current program<br>(mean ± SD) | 3.5 ± 1.8 | 87 |
| % non-university teaching<br>experience | 20% | 93 |
| % other university teaching<br>experience (non-GTA) | 33% | 93 |
| % prior GTA experience | 66% | 73 |
| of which terms GTA<br>(mean ± SD) | 4.1 ± 4.2 | 100 |
| % prior Bis2A GTA<br>experience | 40% | 100 |
| of which terms Bis2A GTA<br>(mean ± SD) | 3.3 ± 2.3 | 86 |

### Training program

Each of the 15 GTAs taught three two-hour discussion sections per week of ~24 students each. Prior to the first discussion section each week, GTAs participated in a two-hour practice-based training session. Trainings were designed to quickly ramp up GTAs’ ability to effectively and consistently implement instructional techniques that emphasized student accountability, logic-development, and practice of problem-solving skills. The training period was divided into two one-hour sessions. The first hour focused on course-specific content review. Content review was conducted as a “mock warm-up”, with training leaders modeling target techniques and GTAs acting as students. During the first five weeks, the second hour of training was dedicated to practice of a specific instructional technique (see “Target techniques and drills” for more details). Prior to GTA practice (“drill”), the theory behind the target technique was explained and step-by-step implementation guidelines were provided, followed by a demonstration by the training leaders. For most drills, GTAs were split into two groups of approximately seven GTAs plus a training leader, with each GTA practicing the target technique under the training leader’s guidance. In week six, the structure of the second hour of training changed and GTAs used this time to collaboratively design interactive, question-based warm-up sessions for their discussion sections using the techniques learned in the first five weeks. The schedule of the training program is shown in **Table 2**. In addition to weekly training, GTAs also attended a one-hour meeting the week prior to the start of term, covering course organization, grading policies, and an overview of the goals of the training program. Undergraduate learning assistants (ULAs) were present during GTA training sessions, but for the most part participated in drills as mock students, not as mock instructors due to time constraints. ULAs served a supplementary instructional role during discussion sessions. However, GTAs did occasionally arrange for a ULA to lead a class session. ULA technique use was included in analysis of student learning outcomes, but not in analysis of GTA teaching practices.

**Table 2:** Training schedule. Focus areas for first and second hours of two-hour weekly GTA training meetings.

| Week | 1st hour | 2nd hour |
| --- | --- | --- |
| 0 | Administration/Buy-in and pedagogy discussion | N/A |
| 1 | Content review | Drill (Cold Call) |
| 2 | Content review | Drill (Stretch It: Explain Logic) |
| 3 | Content review | Drill (Right is Right) |
| 4 | Content review | Drill (Stretch It: Follow-up) |
| 5 | Content review | Drill (Circulate/Check for Understanding) |
| 6 | Content review | Warm-up development |
| 7 | Content review/<br>Drill (Circulate/Check for Understanding) | Warm-up development |
| 8 | No meeting | No meeting |
| 9 | Content review | Warm-up development |
| 10 | Content review | Warm-up development |

### Target techniques and drills

Target techniques were selected that facilitated our overall goal of creating a highly engaged classroom characterized by a high degree of student participation in iterative practice and feedback within a safe and supportive classroom environment. At the beginning of the training program, GTAs were provided with literature describing the target techniques, including the theory behind each technique, implementation tips and selected references to relevant research literature (**Supplementary Document 1**). Target techniques were (in alphabetical order): Circulate/Check for Understanding, Cold Call, Debrief, No Apology, Normalize Error, Praise Effort, Praise Improvement, Right-is-Right, Stretch it: Explain Logic and Stretch it: Follow-up (Lemov 2010). Descriptions and selected references are provided in **Table 3**. Due to time constraints, only five of the ten target techniques were explicitly drilled in training. Drilled techniques were selected from target techniques based on the trainers’ perception of the most urgent needs. All ten techniques were discussed in training sessions and/or one-on-one feedback meetings between the GTA and the training leaders.

**Table 3:**
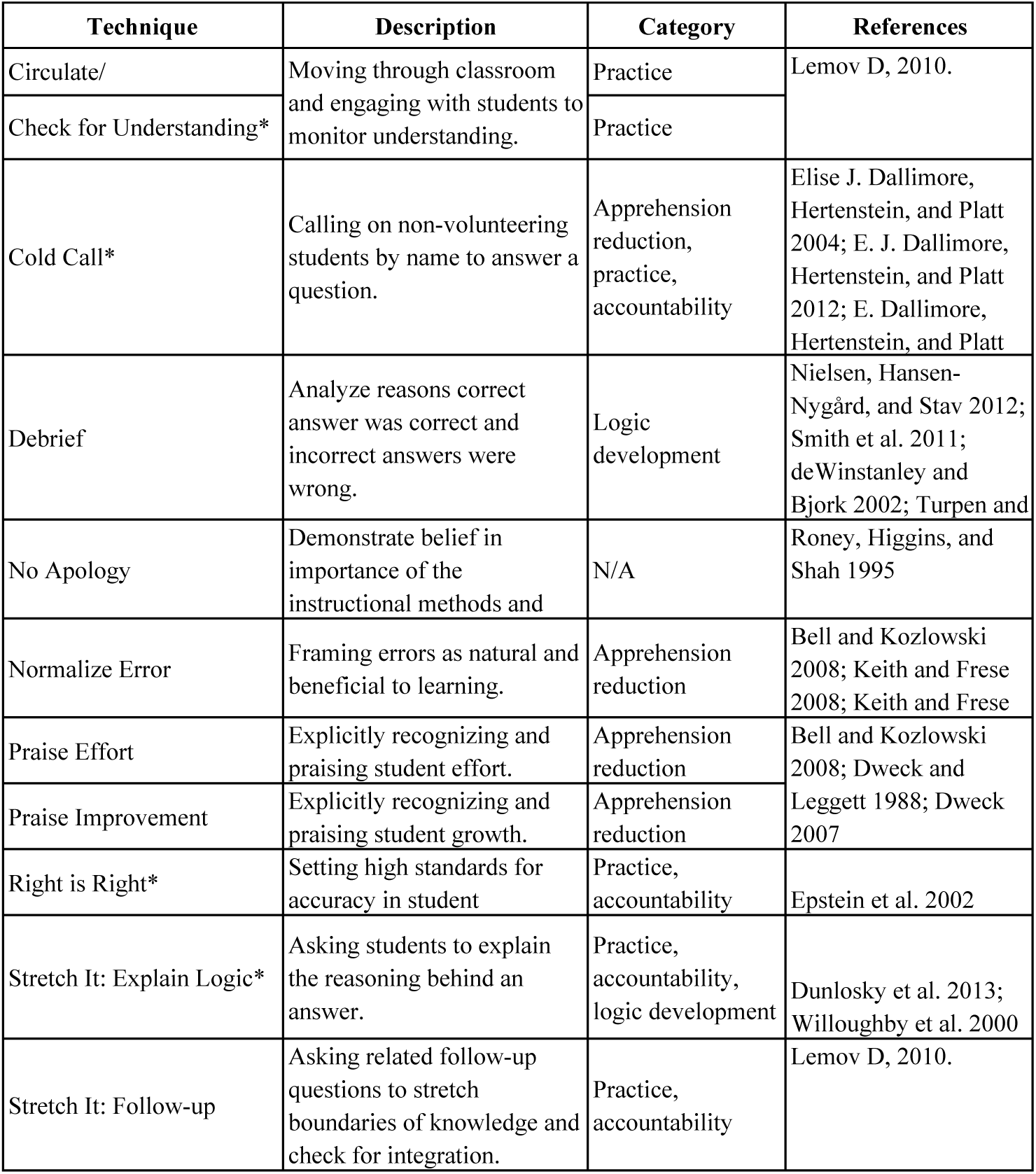
Target techniques. Technique names and descriptions are derived from (Lemov 2010). Selected references are given; however, the same or very similar teaching practices may be referred to by various names in the literature. Techniques marked with an asterisk were drilled during training sessions.

Drills were designed to allow GTAs to practice techniques in a highly simplified mock classroom environment in order to make the techniques routine and habitual. GTAs were informed throughout training that drills were not meant to be accurate representations of authentic classroom experiences. Complications arising in the GTAs’ classrooms and advice for applying the techniques to specific classroom situations were discussed throughout the quarter. After techniques had been explained and modeled by the training leaders, drills were conducted as follows:

#### Cold Call

GTAs were provided with a topic and given three minutes to develop questions on that topic. Each GTA then had three minutes to cold call the other members of their group, with the goal of calling on as many people as possible. GTAs playing the role of students were instructed to always provide correct and complete answers.

#### Stretch It: Explain Logic

Same as the Cold Call drill, with the addition that for each answer provided, the GTA asked the “student” to explain their reasoning before moving to a new question.

#### Right is Right

Same as the Cold Call drill, with the exception that specific “students” were selected to provide incorrect or incomplete answers. The GTA doing the drill was unaware of which “students” would give faulty answers. Upon detecting an incorrect or incomplete answer, the GTA was instructed to restate the question in a way that helped the “student” identify their error. “Students” were instructed to give the correct answer after redirection.

#### Stretch It: Follow-up

Same as the Cold Call drill, with the addition that each “student” called on was asked two to three questions of increasing difficulty on the same topic to check for depth of understanding.

#### Circulate/Check for Understanding

Prior to the drill, training leaders collaboratively identified GTAs needing additional skill development (“rookies”) as well as GTAs with strongly developed skill in this technique (“peer modelers”), based on classroom observations. Each training leader formed a drill team composed of two rookies and one peer modeler. GTAs were not informed if they were a rookie or a peer modeler. While the remaining GTAs worked in small groups to solve mock warm-up problems, the drill teams circulated. First, the training leader stopped at a group and demonstrated proper circulation technique: entry into the group’s conversation, asking for an explanation of the group’s solution or thought process, and ensuring verbal participation from all group members. Each GTA completed this process with different drill groups.

During drill sessions, the training leader and other GTAs provided short (10-15 second) positively-framed feedback for each GTA. Public feedback focused on positive aspects of individual performance, while targeted corrective feedback was reserved for regularly scheduled private feedback meetings (see “Observations and feedback”).

### Observations and feedback

Due to the reported importance of timely, goal-directed feedback in influencing teaching practices (Rezler and Anderson 1971; O’Reilly and Renzaglia 1994; Fedor and Buckley 1987; Gormally, Evans, and Brickman 2014), we incorporated three cycles of feedback into our training program. In-person classroom observations were conducted for each GTA on the second, fourth and seventh weeks of the course. The goal of these observations was to monitor adoption of target techniques and provide frequent, nearterm, and goal-oriented feedback to the GTAs (Rezler and Anderson 1971; O’Reilly and Renzaglia 1994; Fedor and Buckley 1987). In order to ensure that performance goals were clear, feedback focused on a recognized task standard (Hattie and Timperley 2007) using an in-house classroom observation protocol (**Supplementary Document 2**), which was provided to GTAs at the beginning of the term. This protocol also facilitated the development of quantitative personalized improvement goals for each GTA. Upcoming observations were announced in the weekly training meeting, at which point each GTA completed a written self-evaluation (**Supplementary Document 3**), which asked them to assess their own strengths and weaknesses in implementing the target techniques. Weaknesses were phrased as “instructional techniques that you would like to improve” and it was explained that personalized goals would be set during follow-up meetings. Individualized goal setting is thought to improve motivation, leading to increased effort towards the desired task (Gormally, Evans, and Brickman 2014).

Each GTA was responsible for three discussion sections. In-person observations were conducted during the second of each GTA’s discussion sections, allowing the GTAs to practice the techniques unobserved in their first discussion section. The observation period was shortened as the course progressed, with median observation length of 40 minutes in week two, 30 minutes in week four, and 15 minutes in week seven. This decrease in the length of observation was intentional, as observers needed less time to assess classroom practice after becoming familiar with each GTA’s basic teaching habits. Observations were conducted such that each GTA was observed at least once by each of the two training leaders (authors EAB and EJE). As necessary, observations were also completed by a third observer (author CP) who was part of the study design and had been trained on the observation protocol.

At the end of each observation week, the observers met to discuss GTA progress and come to consensus on what personal feedback should be provided to each GTA. This feedback included a written evaluation followed by a face-to-face meeting. The written evaluation included 1-4 observed strengths and 1-3 areas to improve, paired with specific, quantitative goals and was sent to each GTA by email. Each GTA then attended a 15-minute individual face-to-face meeting with one of the observers to discuss their feedback and any issues they were facing in their classrooms. Written summaries for the second and third iterations of the feedback process included whether the previous goal(s) had been met. For the third iteration, written summaries were more informal, representing a summary of the GTAs progress over the quarter, and in-person feedback meetings were not held. Instead, open office hours for discussing feedback were offered; however, none of the GTAs attended these non-mandatory meetings.

### Video coding

To assess the frequency of GTA implementation of target techniques and how GTAs elicited student participation across classrooms throughout the quarter, video recordings were taken of all 45 classrooms in the second, fourth, seventh and tenth weeks of the course. A total of 160 (89%) of the targeted classroom sessions were successfully recorded. Eighty-six percent of missing data (19 videos) were from week seven. All GTAs were recorded between eight and twelve times, with 80% of the GTAs being recorded ten or more times. Data from 159 videos were included in analysis of student learning outcomes; one video was excluded where one GTA substituted for another. Two of the recorded classroom sessions were led by undergraduate learning assistants (ULAs), and were excluded from analysis of GTA instructional practices, but included in analysis of student learning outcomes. Only the first hour of class time was coded, roughly corresponding to the GTA’s individually designed interactive questions and answer (“warm-up”) sessions. In five instances, observations began after the start of class, with a median start time of 53 minutes into class. In three instances, less than one hour of video was recorded, with a mean recorded time for incomplete videos of 52 minutes.

Videos were coded using a modified version of the in-house classroom observation protocol, along with a coding manual (**Supplementary Documents 4 and 5**). Video coding was done by three trained undergraduate observers. Initial training consisted of two one hour-long sessions in which codes were defined, potential scenarios discussed, and short clips of observation videos coded to consensus. All videos were coded using one of four progressive strategies. Initially, observers coded videos in tandem with a partner to provide opportunities to discuss any discrepancies in their understanding of the codes (“paired-tandem”). After solidifying understanding of the codes, observer pairs coded independently and compared results before entering the data in order to resolve major discrepancies (“paired-checked”). After developing expertise in coding, observer pairs coded independently of one another and did not compare results (“paired-independent”). Finally, after confirming high levels of inter-rater reliability (IRR) (see below), individual observers coded the remaining videos independently (“single”). Videos were randomly assigned to the undergraduate observers and observer pairs were rotated throughout the first three stages.

Inter-rater reliability (IRR) was assessed separately for paired-checked (23 videos) and paired-independent (19 videos) video sets. For both sets, IRR was assessed independently for each code using a one-way mixed model, absolute agreement, single-measures interclass correlation (McGraw and Wong 1996; Shrout and Fleiss 1979) to assess the degree of consistency in observer ratings of classroom practices. Inter-class correlation coefficient (ICC) calculations were carried out using the irr R package (Gamer, Lemon, and Puspendra Singh 2015). The resulting ICC values are provided in **Supplementary Table 2**.

Only codes with an ICC value >0.75 (“excellent”) in the paired-independent video set were included in further analyses. High ICC indicates that these classroom practices and participation types were rated with a high degree of fidelity across observers, suggesting that a low degree of measurement error was introduced. Consequently, statistical power for further analysis was not significantly impacted by independent coding. These 42 codes were therefore deemed to be suitable to use in further hypothesis testing. Due to this high degree of consistency, the remaining videos were coded by a single observer. Note that ULA classroom practices are included only in analysis of student learning outcomes, and not analysis of GTA practice. Code frequencies were averaged between observers. Codes for class sessions with less than one hour of recorded video were time-normalized to 60 minutes.

Student participation events were coded as one of five types: 1) Student question (student asks a question of the GTA or ULA), 2) Cold Call – Individual (GTA asks an individual student a question without the opportunity to discuss with group members), 3) Cold Call – Group (GTA asks an individual student a question after that student has had the opportunity to discuss the question with group members), 4) Volunteer – Individual (student answers a question without prompting by GTA or ULA), and 5) Volunteer – Group (GTA calls on a group of students to answer a question, of whom one of the students volunteers to provide the answer). In investigating student participation levels, all categories were analyzed independently, as well as in appropriate combinations.

For GTA interactions with small groups (Circulate), a distinction was made between interactions initiated by the GTA (“active” Circulate) and by the students (“moderate” Circulate). Instances in which the GTA was moving throughout the classroom but not interacting with students were captured by the “passive” Circulate code. For more details on code definitions, see **Supplementary Document 5**.

### Longitudinal analyses of classroom practice

To understand the dynamics of GTA instructional practices, we analyzed changes in frequency of technique use throughout the quarter, as well as changes in student participation in class discussions. For each week, the frequency of each coded technique or participation type was averaged for each GTA across their recorded sessions (usually three). For each technique, we investigated whether there were differences in GTA practice between the beginning (week two) and end (week ten) of the course. Since the frequency for most techniques did not meet assumption of normality, statistical significance of difference in means was tested using the non-parametric Wilcoxon rank-sum test in R (R Core Team 2014). Effect sizes were calculated using Cliff’s d, which is an ordinal statistic describing the frequency with which an observation from one group is higher than an observation from another group compared with the reverse situation (Cliff 1993). Cliff’s *d* can be interpreted as the degree to which two distributions (x and y) overlap, with values ranging from -1 to 1. A Cliff’s *d* value of 0 represents no difference in the sample distributions, a Cliff’s *d* value of −1 indicates that all samples in distribution x are lower than all samples in distribution y, and a Cliff’s *d* value of 1 indicates the opposite. Threshold values for Cliff’s *d* used throughout are defined in (Romano et al. 2006) as implemented in (Torchiano 2015). This method has been shown to be quite robust to violations of normality and heterogeneity of variance (Cliff 1993). Cliff’s *d* calculations were done using the effsize R package (Torchiano 2015).

### Analysis of feedback

Trainers met with each GTA the week after their classrooms were observed to discuss strong points and potential areas of improvement in classroom management, technique implementation and content knowledge. Each GTA was also given a written summary of feedback prior to the in-person meeting. The feedback given to the GTAs was characterized independently by the training leaders (EAB and EJE), who analyzed the written feedback summaries for each GTA each week and coded them for presence or absence of appreciation feedback (in which the GTA was praised or acknowledged for proper technique use or high technique frequency) or coaching feedback (in which GTA was prompted to improve fidelity or frequency of technique implementation) (Stone and Heen 2014) for each of the target techniques. Discrepancies in independently derived codes were discussed by the training leaders until consensus was reached. For each technique, each GTA was coded as having received neither type of feedback, appreciation only, coaching only, or (in rare cases) both types of feedback, for each week in which feedback was given.

For each technique, the change in technique frequency between the observation immediately preceding feedback and immediately following feedback was compared between 1) GTAs receiving appreciation feedback, 2) those receiving coaching feedback, and 3) GTAs receiving no feedback for that technique. Change in technique frequency, rather than absolute frequency, was used as an outcome metric due to longitudinal differences in technique implementation. Statistical significance of difference in means was tested using the non-parametric Wilcoxon rank-sum test, and effect sizes were calculated as Cliff’s *d* (Cliff 1993), using the effsize R package (Torchiano 2015).

### Relationship between classroom practice and student learning outcomes

We investigated the relationship between GTA/ULA classroom practices and student learning outcomes using multiple linear regression with overall course exam points as the response variable. We used exam points, rather than GTA-awarded points, to avoid confounding GTA teaching effectiveness with GTA grading leniency. As discussion enrollment was not randomized, student demographics were incorporated into statistical modeling as potential confounding variables for their success in the course. The importance of correcting for differential student demographics in non-randomized educational studies has been demonstrated by Theobald and Freeman (2014).

To determine appropriate student demographic variables for inclusion in the model, we used the previous and subsequent Fall terms (Fall 2013 and Fall 2015) as training data sets. These terms were taught by the same lecture instructor as the study term (Fall 2014). First, we established that prior academic achievement (cumulative institutional GPA) and demographics (gender, transfer status, first generational student status, under-represented minority (URM) status, and course repeater status) were similar across all three terms (**Supplementary Table 3**). To enable inclusion of new transfer students and freshmen, cumulative institutional GPA used was that from the end of, rather than beginning of, the term. Grade earned for the study course was excluded from the calculated GPA. First generation students were defined as students whose parent(s) or legal guardian(s) had not completed a bachelor’s degree. URM was defined as anyone who self-identified ethnically as African-American/Black, Puerto Rican, American Indian/Alaskan Native, Mexican-American/Mexican/Chicano, Latino/Other Spanish, or Hispanic-Other. Transfer students were defined as anyone who had transferred to the university from an accredited community college. Demographic data was compiled from centralized university databases.

Model selection for the two training datasets was automated using the cross-validate model selection function in the glmnet R package (Hastie and Qian 2014). For each training data set, the selected model was that which had the lowest mean-squared error within one standard deviation of the minimum value for the regularization parameter lambda. Both models included GPA, gender, and first generational status but did not include students’ URM or repeater status. The model for F15 also included students’ transfer status. We used these models as a starting point for developing an appropriate model for the study term.

To model student learning outcomes as a function of GTA behavior, we employed a data reduction technique, reducing all coded behaviors to a series of aggregate variables. The technique used was principle component analysis (PCA), a data reduction technique for summarizing the information contained in several variables (in our case, coded behaviors), via a smaller number of aggregate variables (Cudeck 2000). A total of nine coded behaviors [Cold Call rate (%), volunteer rate (%), student question rate (%), Circulate (passive), Circulate (moderate), Circulate (active), Right is Right, Stretch-it: Explain Logic, and Stretch-it: Follow-up] were used in the final PCA. Statistically significant outliers (assessed using Mahalanobis distance (Mahalanobis 1930), cutoff chi-square > 20) were removed from the dataset to yield a final analytical sample of nine coded GTA/ULA behaviors and 132 total classroom observations. Note that, although a PCA analysis was conducted, due to the relatively small sample size of our observations (n = 132), a statistical reduction technique such as PCA may not be appropriate and may lead to inaccurate component estimations. PCA was conducted only as an exploratory analysis to see which potential components could be extracted.

PCA analysis was run using the psych R package (Revelle 2015). An oblimin rotation was used to extract a total of two components. An oblimin rotation was selected to allow the extracted components to have some potential correlation between them (Kim and Mueller 1978). The two components were named “accountability” and “volunteer rate” and together accounted for 48% of the observed variation in classroom practice. Accountability included Cold Call rate (%), Right is Right, Stretch-it: Explain Logic, and Stretch-it: Follow-up. Volunteer rate was the percent of unique students who volunteered to answer a question. To create an aggregate variable for accountability, the individual variables for that component were normalized (mean = 0, SD = 1) and then averaged, creating a z-score for use in the regression model.

Based on models recovered from the training data sets, and components recovered from PCA analysis, the starting model for the study term was:

Total_exam_points ~ cumulative_GPA + gender + first_generation_status + accountability + percent_volunteer.

Accountability and percent volunteer metrics were constant for all students in a particular discussion section, while other variables represent values for individual students. To this base model, we tested addition of students’ transfer status, GTAs’ experience level (number of times during study term GTA had taught that discussion material prior to the discussion in question) and GTA identity (i.e. which GTA taught the section). GTA experience and identity were individually added as random effects using the nlme R package (Pinheiro et al. 2014) and improvement to model was tested by ANOVA. Neither significantly improved the model (cutoff ANOVA *p* < 0.05). Student transfer status was added as a fixed effect and tested by ANOVA. Although addition of transfer status significantly improved the model (*p* = 0.030), adjusted R^2^ increased only marginally (Δ = 0.0026) and therefore this term was not incorporated into model. The final model was thus identical to the starting model described above.

Students missing demographic information for variables included in final model were excluded from analysis (n = 31 students, 3.1%). Students who did not take all required examinations were also excluded (n = 13 students, 1.3%). A total of 946 students (95.5%) were included in the final analysis.

The final model was tested for conformity to assumptions of linear modeling: normality of residuals (by Shapiro-Wilk test (R Core Team 2014), cut-off value W > 0.95 or *p* > 0.05), constant variance (by Breusch-Pagan test (bptest in R package lmtest (Hothorn et al. 2011), cut-off value *p* > 0.05), and lack of overly influential data points (by Cook’s distance (cooks.distance (R Core Team 2014), cut-off value 1). As the model displayed heteroskedasticity, a heteroskedasticity-corrected covariance matrix was calculated using hccm in the R car package (Fox and Weisberg 2011), with the classical White correction. This corrects standard error estimates to account for heteroskedasticity, but does not affect model coefficients.

### GTA exit survey

In the final week of class, an online survey was circulated to the GTAs to assess their opinions about the training program and the class as a whole (**Supplementary Document 6**). The survey was divided into three sections. Section A (six questions) asked GTAs to rate the usefulness of components of the training program (including feedback given, drills, and content review) on a five-point Likert scale ranging from 5=very helpful to 1=very unhelpful. Section B (18 questions) asked GTAs to rate statements about the clarity of course expectations, positivity of interactions with different course participants, positivity of the overall GTA experience, self-assessed improvement of their teaching abilities and usefulness of the target techniques on a five-point Likert scale ranging from 5=strongly agree to 1=strongly disagree. An N/A option was also provided. Section C (four questions) consisted of free-response questions regarding future improvements to GTA training, their overall course experience (both positive and negative), and general comments. Eleven of fifteen GTAs (73%) completed the survey.

### IRB

This study was deemed exempt from full IRB review and determined to not be research involving human subjects as defined by the Department of Health and Human Services. IRB ID #513796-1.

## Results

### GTA classroom practices and student participation levels

We first sought to describe overall GTA classroom practice in terms of frequency of utilization of the target techniques and levels of student participation. Although each of the target techniques listed in **Table 3** was included in the GTA information packet at the start of the term and discussed in training sessions, only five techniques were drilled (Circulate, Cold Call, Right is Right, Stretch it: Explain Logic and Stretch it: Follow-up). Significantly, techniques that were not drilled occurred rarely, at frequencies too low to pass our filter for inter-rater reliability (IRR). Overall frequency (across all 158 classroom observations) for the ten target techniques is given in **Table 4**. Five number summaries for each of the drilled techniques are given in **Supplementary Table 4**.

**Table 4:**
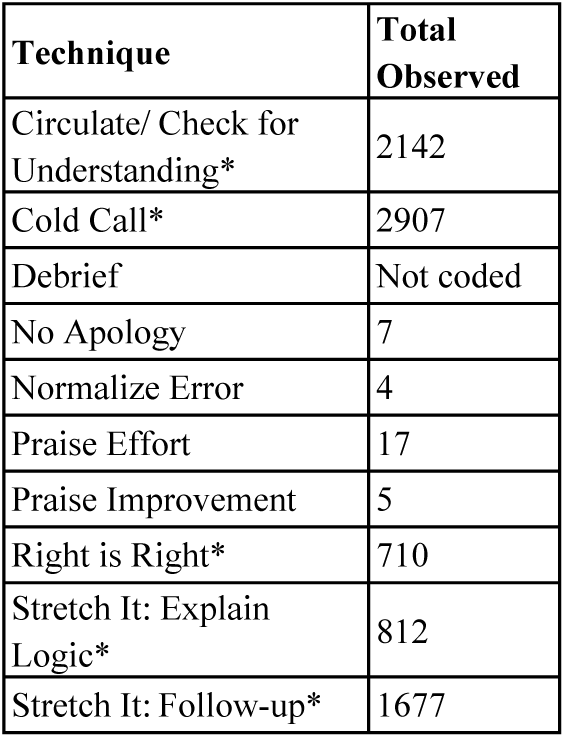
Target technique total observed frequency. Only GTA activities (not ULA) are included in counts. Passive Circulation (in which GTA moved throughout the room, but did not interact with students) is excluded. The Debrief technique was not coded due to inability to reach consensus coding criteria. Counts represent raw sums from all 158 classroom sessions in which a GTA acted as the primary instructor (excluding two observations in which a ULA acted as instructor). Techniques marked with an asterisk were drilled during training sessions.

Cold Call was the most frequently utilized technique (**Supplementary Table 4**), with a median frequency of 18 events per hour observed. Circulate was the next most common, at 13 events per hour when both moderate (student-initiated) and active (GTA-initiated contact) were considered. The other techniques which were used often enough to quantify accurately (Right is Right, Stretch it: Follow-up, and Stretch it: Explain Logic), were implemented with lower frequencies. This was not surprising, given that these techniques, by definition (see **Supplementary Document 5**), could only follow a Cold Call event, and could not be initiated independently. When occurrence of these three techniques are summed, their frequency is similar to Cold Call (median of 19.73 compared with 18.17, *p*-value of difference between means = 0.81) with strong positive correlations between the number of Right is Right, Stretch it: Follow-up and Stretch it: Explain Logic events and the percent of students in a classroom who were cold called (**Supplementary Figure 1**). This relationship indicates that GTAs using Cold Call were doing so in conjunction with additional techniques designed to improve students’ accountability for their knowledge and provide opportunities for practice and logic development.

In investigating how participation was manifested in our active learning classrooms (i.e. which types of participation were present and what proportion of students participated), we sought to answer the intertwined questions of 1) which of the four participation mechanisms we measured (individual volunteer, group volunteer, individual cold call and group cold call) were most successful in prompting high proportions of unique students to respond and 2) whether the more successful mechanisms were preferentially utilized by GTAs. These questions were addressed in order to determine whether GTAs were making optimal use of these techniques for eliciting student participation.

First, for the two response mechanisms for which GTAs called on individual students, we investigated whether GTAs were more likely to call on unique students without providing time for students to discuss the problem with their groups (individual cold call) or after such a discussion (group cold call). Although the percentage of unique responders was high for each response type (mean of 81% and 89% respectively), GTAs were significantly more likely to call on new students after group work than when no group work time was provided (Cliff’s d = −0.41, 95% CI [−0.6, −0.18], *p* = 8×10^−4^) (**Figure 1A**). Despite being more effective in producing unique participants, the group cold call participation mechanism was used less frequently by GTAs than individual cold call (Cliff’s d = 0.54, 95% CI [0.33, 0.71], *p* = 9.2×10^−6^) (**Figure 1B).**

**Figure 1:**
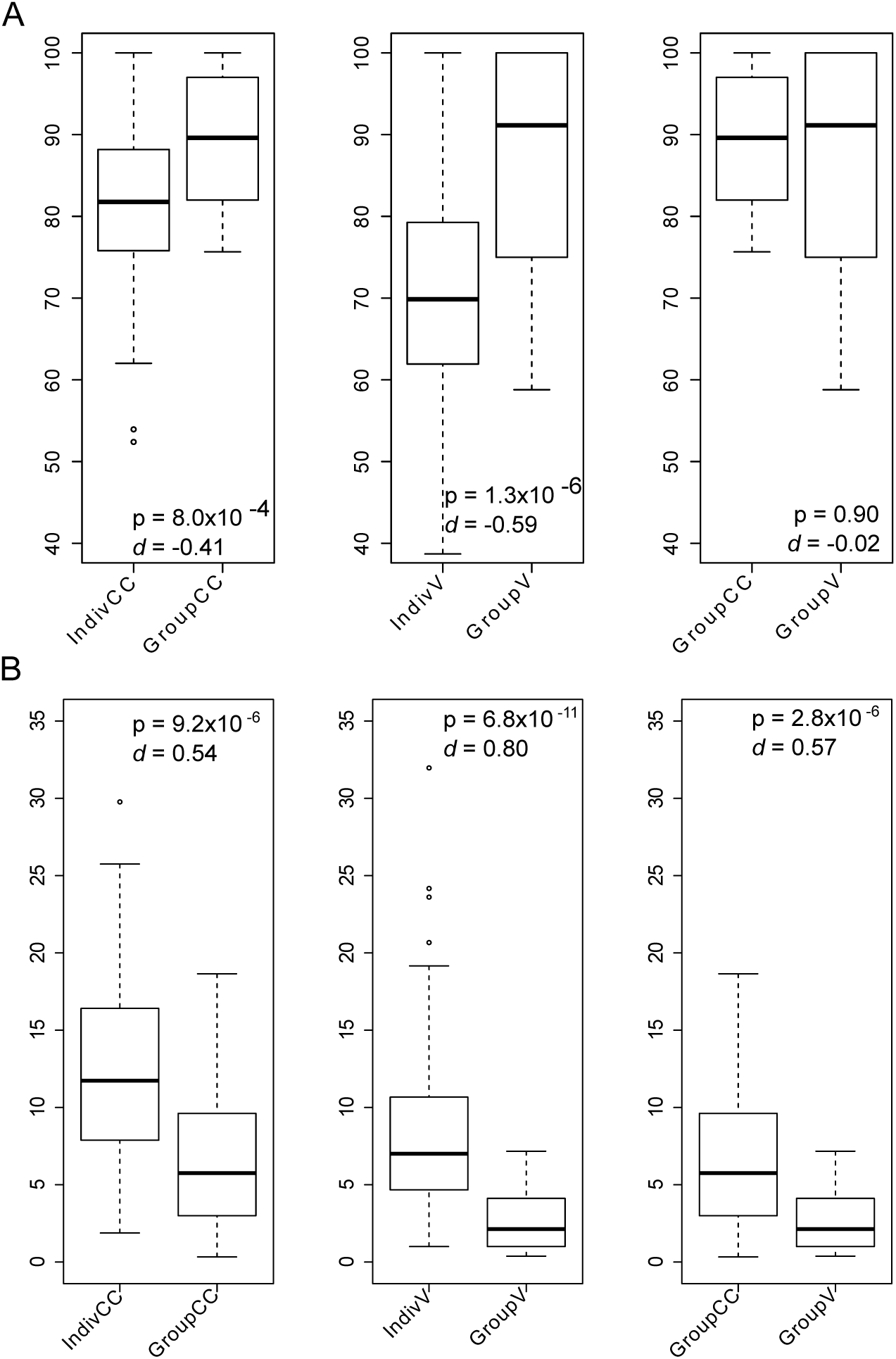
Participation types and unique responder rates. (A) Percent of responders who were unique for each type of participation. (B) Frequency of different participation types. All values are averaged across observed classroom sessions for each individual section (n = 45 sections). IndivCC = Individual Cold Call, GroupCC = Group Cold Call, IndivV = Individual Volunteer, GroupV = Group Volunteer. Negative values of Cliff’s *d* indicate technique on the right of each pair occurs at higher frequency (panel B) or has higher proportions of unique responders (panel A). Positive values indicate the opposite. For more information about interpreting Cliff’s *d* values, see Methods.

Next we asked whether GTA-targeting of specific groups to answer a question (group volunteer) led to higher levels of unique participants than relying on individual students to volunteer. Individual volunteers showed a strong bias towards repeated response from the same students (mean unique responder rate of 70%) compared with GTA-targeted procurement of volunteers from a specific group (mean of 87% unique) (Cliff’s d = -0.59, 95% CI [−0.74, −0.38], *p* = 1.3×10^−6^) (**Figure 1A**). Although calling on groups led to a more diverse set of participants, GTAs were significantly more likely to take volunteer responses from the class as a whole than to request an answer from a particular group (Cliff’s d = 0.80, 95% CI [0.64, 0.89], *p* = 6.8×10^−11^) (**Figure 1B**).

Finally, for instances in which students were asked to respond to questions following group work, we asked which of two strategies - cold calling an individual student, or asking for volunteers from the group - was likely to lead to a greater proportion of unique responders. We found no significant difference in unique responder rate between the two strategies (Cliff’s d = −0.02, 95% CI [−0.26, 0.23], *p* = 0.89), although GTAs were more likely to ask for volunteers than to cold call students after group work (Cliff’s d = 0.57, 95% CI [0.35, 0.74], *p* = 2.6×10^−6^).

In addition to counting frequency of implementation for each of the ten target techniques, we were also interested in measuring the fraction of students in each classroom who were given the opportunity to participate in the whole-class discussion. Three-quarters of classrooms had an average participation level above 70% (range: 29%-100%) and a Cold Call participation level above 46% (range 14%-100%) (**Table 5**). When GTAs are considered as the unit of interest, three-quarters averaged overall participation levels above 64% and a Cold Call participation level above 50%. In the median classroom, only 15% of students asked a question during whole-class discussion (range: 3%-37%), highlighting the inadequacy of relying upon spontaneous student questions for gauging understanding. See **Supplementary Table 5** for breakdown of participation types. For this analysis, we were interested in GTA attempts to implement the target techniques; therefore, here we included students who were called on to answer a question, but did not respond. Overall non-response to Cold Call was 4.4%. Note that this measure does not capture student-student interactions in small groups, nor interactions between GTAs and students during small-group work; the later factor is captured in the Circulate (active) and Circulate (moderate) codes.

**Table 5:**
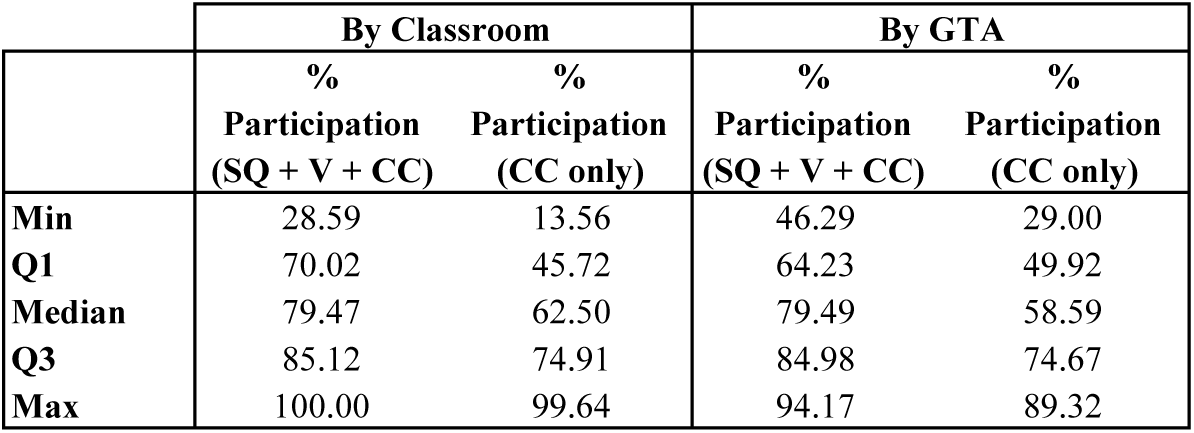
Participation statistics. Summary statistics for participation levels averaged across all observed classroom sessions for each classroom (left) or each GTA (right). Numbers represent percent of students in class on the day observed who participated in whole-class discussion. Overall levels of participation (including student questions (SQ), volunteer responses (V) and Cold Call (CC)), and Cold Call levels separately are shown. For more detailed breakdown of participation types, see **Supplementary Table 5**. N = 45 classrooms, 15 GTAs.

|  | By Classroom |  | By GTA |  |
| --- | --- | --- | --- | --- |
|  | %<br>Participation<br>(SQ + V + CC) | %<br>Participation<br>(CC only) | %<br>Participation<br>(SQ + V + CC) | %<br>Participation<br>(CC only) |
| **Min** | 28.59 | 13.56 | 46.29 | 29.00 |
| **Q1** | 70.02 | 45.72 | 64.23 | 49.92 |
| **Median** | 79.47 | 62.50 | 79.49 | 58.59 |
| **Q3** | 85.12 | 74.91 | 84.98 | 74.67 |
| **Max** | 100.00 | 99.64 | 94.17 | 89.32 |

### Changes in classroom practice over course of training program

To characterize possible impacts of the training program, we investigated whether GTA classroom practices changed over the course of the term. Of the 16 techniques and participation types measured, six showed significant changes between the first and last observation periods (**Figure 2** and **Supplementary Table 6**). Frequency of Cold Call, and Stretch it: Explain Logic significantly decreased over the course of the term, as did overall participation rate and Cold Call levels. In contrast, the frequency of student-initiated contact with the GTA during small-group work (Circulate (moderate)) increased, as did the frequency of Group Volunteer events (in which GTA called on a specific group, but allowed students within that group to decide who would answer the question).

**Figure 2:**
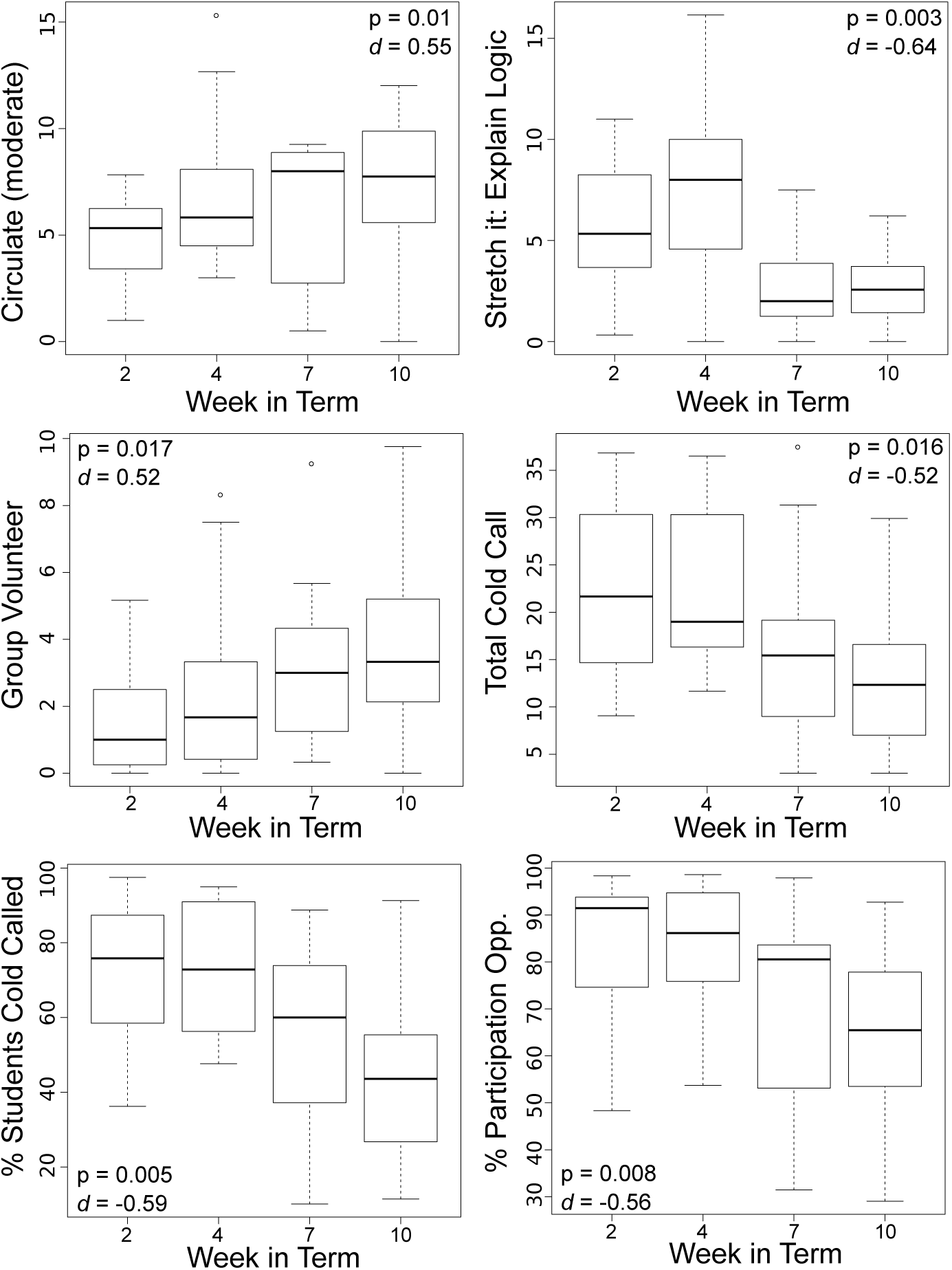
Longitudinal changes in GTA classroom practices. Boxplots showing distribution of technique and participation type frequency for each week in which classroom observations were done, averaged across all observed sections for each GTA (usually three). Box sections represent second and third quartiles. Whiskers represent first and fourth quartiles. Thick line represents medians. Outliers are shown with open circles. P-values and Cliff’s *d* values are for difference between weeks 2 and 10. Negative values of Cliff’s *d* indicate decrease in frequency of practice, positive values indicate increase in frequency. For more information about interpreting Cliff’s *d* values, see Methods. N = 15 each for weeks 2, 4 and 10 and 11 for week 7. Boxplots for techniques and participation types not shown here are available in **Supplementary Document 7**. See **Supplementary Table 6** for all longitudinal comparisons.

### Changes in classroom practice following feedback

To explore the potential impact of technique-specific feedback on GTA practice, we looked at changes in technique utilization between the observation immediately prior to and immediately after each feedback session. We compared the change in use of specific techniques for GTAs who received either coaching feedback or appreciation feedback against those who received no feedback for that technique. The Stretch it: Explain Logic technique was not included in this analysis, due to a difference in how the two variants of this technique were coded, preventing the pooling of observations for instances in which the same or a different student was asked to provide an explanation (see **Supplementary Document 5**). Circulation (moderate) (i.e. student-initiated GTA-small group contact) was also excluded, because GTAs had no direct control over how frequently this type of interaction occurred. As discussed previously, the No Apology, Normalize Error, Praise Effort, and Praise Improvement techniques were not used often enough to accurately quantify. Thus, only four of the ten target techniques were included in analysis of feedback impact.

Of these four, a significant positive relationship between coaching feedback and technique frequency was found for Right is Right (*p* = 0.04) and Circulate (active) (i.e. GTA-initiated small-group contact) (*p* = 0.03) (**Figure 3**), indicating that GTAs receiving coaching feedback for these techniques subsequently increased their implementation frequency compared with GTAs who received no feedback on these techniques. Coaching feedback was only provided once for Stretch it: Explain Logic; thus we are unable to make any claims about the effectiveness of this type of feedback for this technique. We also observed a negative relationship between appreciation feedback and technique frequency for Stretch it: Explain Logic (*p* = 0.009). However, no significant relationship was found between appreciation feedback and implementation frequency for other techniques. In contrast to the other three techniques investigated, no significant change in use of Cold Call was observed regardless of the type of feedback provided, suggesting that GTA use of Cold Call was not easily influenced by formal advice.

**Figure 3.**
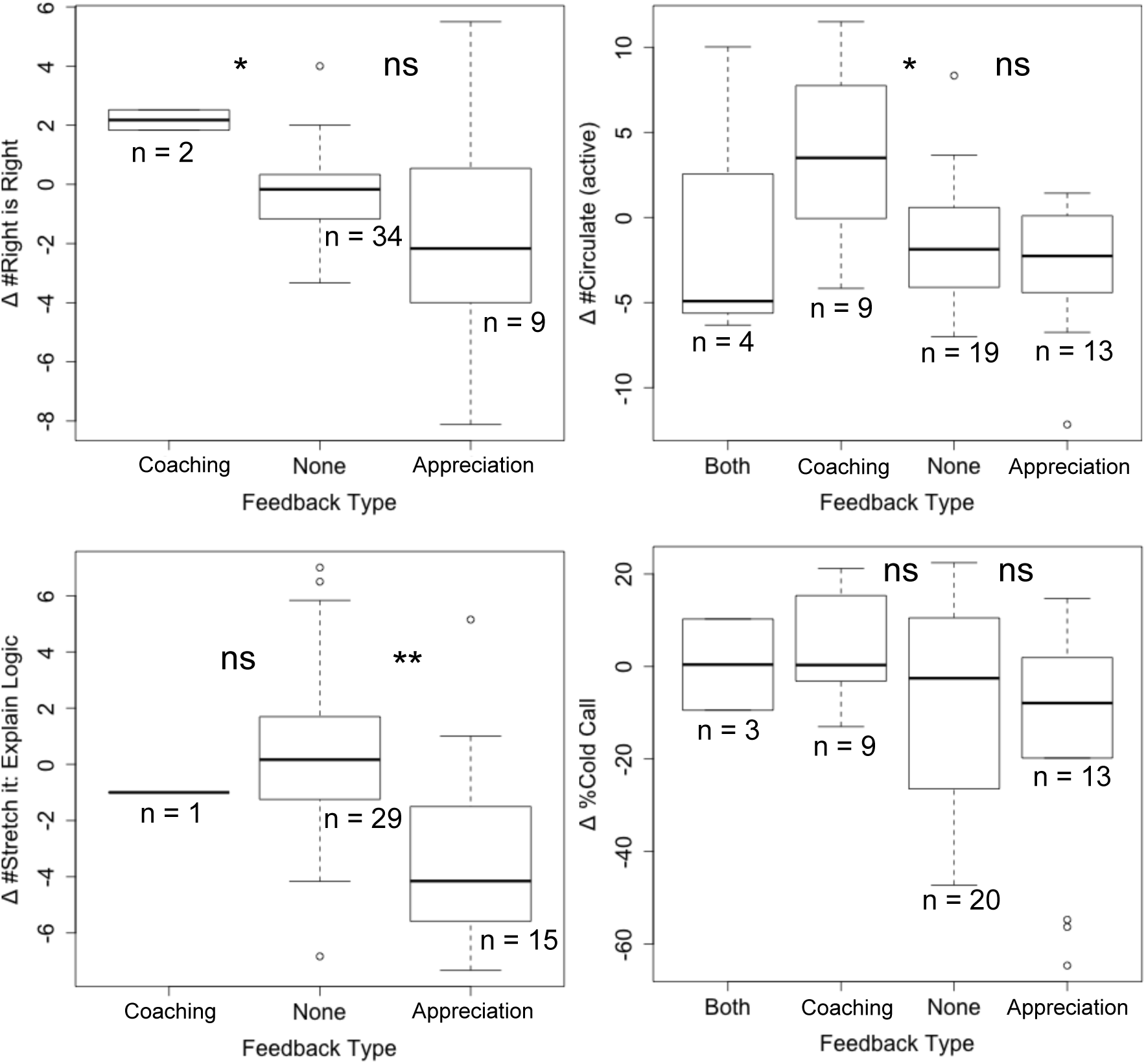
Changes in GTA classroom practices following feedback. Changes in technique frequency between pre-feedback and post-feedback observation sessions. Box sections represent second and third quartiles. Whiskers represent first and fourth quartiles. Thick line represents median. Outliers are shown with open circles. Coaching = feedback intended to bring practice closer to defined task standard, appreciation = feedback that recognizes performance in line with task standard. N = number of instances where particular type of feedback was given. ^*^ = *p*-value < 0.05, ^**^ = *p*-value < 0.01, ns = *p*-value > 0.05. See text for *p*-values.

### Student learning outcomes

In an attempt to determine whether any of the teaching practices we measured were associated with student learning, we modeled the relationship between these practices and student exam scores. Classroom behaviors were condensed into components via principle components analysis. The two components that most fully explained variability in classroom practices were 1) accountability (which included % Cold Call, Right is Right, Stretch it: Explain Logic and Stretch it: Follow-up, see **Table 3**) and 2) % volunteer rate. Although neither of these two components were significantly correlated with student learning outcomes at the *p <* 0.05 confidence level (**Table 6**), the indication of a possible negative trend between student exam scores and volunteer rates warrants further research. We speculate that a negative relationship between the rate of volunteer responses and student performance may exist, due to reduced class-wide attentiveness when volunteers are used as the primary mechanism for eliciting responses. Although our data does not offer strong support for this claim, it suggests that further investigation of this research question may be illuminating. Although the sample size for our study was quite large in the context of classroom observational studies, due to high amounts of natural variation in instructional practices across GTAs, it may be necessary to collect data on an even larger number of class sessions in order to resolve these relationships.

**Table 6:**
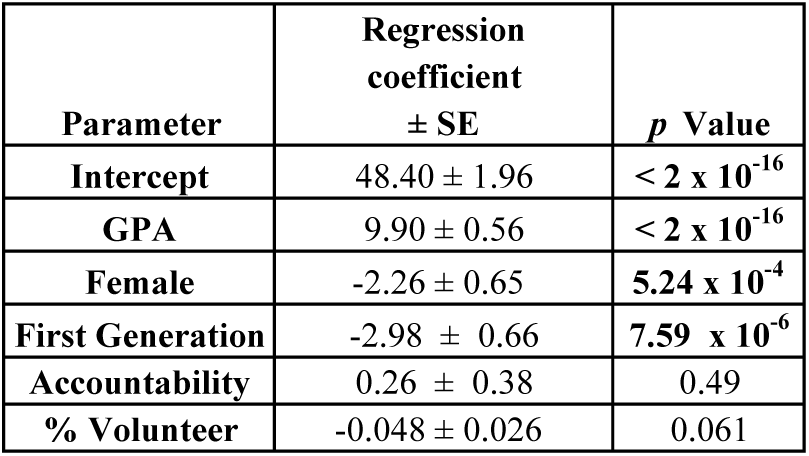
Student learning outcomes. Student exam performance was not significantly associated with either classroom volunteer response levels or accountability (combination of % Cold Call, Right is Right, Stretch it: Explain Logic, and Stretch it: Follow-up). Bolded *p* values are significant at the *p* < 0.05 level. Coefficients are in terms of percentage of exam points.

| Parameter | Regression coefficient<br>± SE | *p* Value |
| --- | --- | --- |
| Intercept | 48.40 ± 1.96 | < 2 x 10<sup>-16</sup> |
| GPA | 9.90 ± 0.56 | < 2 x 10<sup>-16</sup> |
| Female | -2.26 ± 0.65 | **5.24 x 10<sup>-4</sup>** |
| First Generation | -2.98 ± 0.66 | **7.59 x 10<sup>-6</sup>** |
| Accountability | 0.26 ± 0.38 | 0.49 |
| % Volunteer | -0.048 ± 0.026 | 0.061 |

### GTA attitudes

GTA exit survey respondents (11/15, 73%), reported feeling most components of the training program to be useful, including in-person feedback meetings (A.1), feedback summaries delivered by email (A.2), drills (A.4), and time in GTA meetings dedicated to warm-up design (A.5) and content review (A.6). Pre-observation self-assessments were less valued (45% rating neutral or very unhelpful). Three GTAs expressed a desire for more review of course content (B.16) and four reported that the total workload did not meet their expectations (B.17). Overall, GTAs expressed positive attitudes about their involvement in the program, reporting that they felt valued as instructors (B.12), that the techniques covered in training were useful for their students (B.14), and that this experience improved their teaching abilities (B.13). All survey respondents rated their overall experience as a GTA for the course in the study term as positive (B.18). See **Supplementary Table 9**.

## Discussion

Evidence has shown improved student learning outcomes in classroom environments where students take an active role in interacting with the material (Freeman et al. 2014). Effective implementation of evidence-based teaching practices, including active learning, is a highly complex skill requiring deliberate practice to master. Here we sought to transfer a practice-based training framework from its original context in K-12 teacher training to a higher education context, and to demonstrate its ability to help novice higher education instructors (here GTAs) learn and implement evidence-based practices.

Although there are many different aspects of active learning, important components with demonstrated positive effects on student learning have recently been reviewed (Eddy, Converse, and Wenderoth 2015). In this study, we focused on two major hurdles for the implementation of these identified components - expanding the penetrance of active learning-based teaching methods through GTA training, and determining what factors influenced the adoption of these techniques in the classroom. To begin to address these broad goals, we assessed whether a practice and feedback-based training program could be used to train graduate teaching assistants to implement specific active learning practices with high fidelity. We also examined whether student achievement tracked with any of these specific teaching practices as implemented by our GTAs.

We found that, given practice-based training, GTAs were capable of implementing evidence-based teaching practices. However, although initial use of drilled practices was high, adoption was not stable, particularly for participation enforcement techniques (e.g. Cold Call), which decreased over the course of the term. These same techniques were not amenable to formal feedback, as GTAs’ use of these techniques declined despite coaching. Analysis of student participation suggests specific changes in our training methods may ameliorate this effect and increase student engagement. We also found that, although drilled practices were initially adopted with some declining over the course of the term, non-drilled practices were not used with any appreciable frequency, supporting the importance of practice-based training for influencing teaching practices.

GTAs involved in our training program reported positive responses to many aspects of the program, including the use of drills in training sessions and the ways in which feedback was delivered (**Supplementary Table 9**). Utility of the feedback program was further supported by direct evidence of the effectiveness of feedback for at least two of the four assessed techniques (Right is Right and Circulate (active)) (**Figure 3**). These results indicate that at least some of the targeted instructional behaviors were responsive to feedback. However, the lack of any significant change in GTA implementation of Cold Call after feedback indicates that certain instructional practices may be more resistant to external influence. Instructor resistance to Cold Call has been previously documented (E. Dallimore, Hertenstein, and Platt 2006) and matches the authors’ experiences in a variety of instructional coaching settings.

GTAs also reported an overall belief in the usefulness of the target techniques for their students (**Supplementary Table 9**). However, this reported buy-in did not result in consistent levels of some of the targeted classroom practices, specifically Cold Call and the closely-linked Stretch it: Explain Logic technique, which each decreased significantly over the course of the term (**Figure 2**). This result is particularly intriguing given that Cold Call was the only technique for which an explicit performance target (100%) was set. In addition to providing more evidence for the difficulty of influencing instructor use of Cold Call and related accountability techniques, these results highlight the importance of directly assessing classroom practice, rather than relying on self-reported evidence. However, asking GTAs about their perceptions of the utility of each of the individual target techniques may have provided more discrete information and enabled us to more directly assess both buy-in for individual techniques and the extent to which this buy-in translated into adoption of these practices.

The four metrics which decreased over the course of the term were all directly or indirectly tied to Cold Call utilization (**Table 7** and **Figure 2**). Both the total number of times that Cold Call was implemented, and the proportion of students in a classroom who were asked to respond via Cold Call, decreased. One technique which could only be implemented as a follow-up to Cold Call (Stretch it: Explain Logic), also decreased. The decrease in Cold Call and Stretch it: Explain Logic correlated with the cessation of drilling after week five. Although we cannot casually link these events, this fact, in combination with the lack of GTA use of non-drilled techniques, supports the importance of deliberate practice for transforming instructional practices.

This decrease in Cold Call was linked to an overall decrease in participation levels, as, despite the increase in one type of volunteer response (Group Volunteer – GTAs asking for a volunteer respondent from a particular group), overall shifts in volunteer rates were not sufficient to compensate for decreased levels of enforced participation. We did not observe an increase in total volunteer rates after exposure to Cold Call, as had been previously reported (E. J. Dallimore, Hertenstein, and Platt 2012). However, we did find that students became more likely to initiate contact with their GTA during group work time as the term progressed. This result provides some evidence for the idea that students in classrooms with high levels of enforced participation are more likely to take an active role in their education, although the mode of action of this general principle may differ across instructional environments (i.e. increased volunteer rates in (E. J. Dallimore, Hertenstein, and Platt 2012) versus initiating interaction with instructor in this study).

One of the reasons for increasing classroom participation rates was to increase the likelihood for participation from the maximum possible number of students. Therefore, we assessed whether GTAs preferentially used participation mechanisms which were more effective at eliciting unique respondents. We found that GTAs’ preferred implementations of the Cold Call and Volunteer participation types favored individual rather than group responses, despite the fact that the group response mechanism in both cases turned out to be more successful at eliciting unique respondents (**Figure 1**). In terms of concrete classroom practices, this meant that GTAs were more likely to ask questions of students without providing group work time, and that in cases where group work time was provided, GTAs were more likely to ask for volunteers from the room at large rather than from a particular group. We speculate that this first preference may be an artifact of the way drills were conducted, as the drills for Cold Call and related techniques did not involve group work.

We found no indication that calling on individual students following group work (Group Cold Call) was more successful at eliciting unique respondents than calling on groups (Group Volunteer). As the Group Volunteer participation technique was one of only two practices to increase significantly over the course of the term, this result opens up the potential for shifting towards this, perhaps less intimidating, form of participation enforcement. By emphasizing use of Group Volunteer rather than Cold Call in future iterations of the training program, we may be able to improve classroom participation rates, as this engagement mechanism was organically favored by GTAs.

This analysis of GTA use of participation mechanisms suggests two concrete changes to our training program to increase student participation. First, Cold Call drills should provide time for mock students to discuss questions as a group prior to being called on. Second, a variation of the Cold Call drill should be added which prompts instructors to call on groups of students rather than individuals. The data suggest that these two modifications of the Cold Call technique would help overcome instructor resistance to Cold Call, while maximizing unique student participation.

Along with investigating longitudinal changes in GTA instructional practices, we also questioned whether these changes were likely to be attributable to the training program or to organic changes in teaching practices as GTAs became more experienced. The results of our feedback analysis provide evidence that at least a subset of the changes we observed were responses to the feedback component of the training program (**Figure 3**). We therefore propose that many of the longitudinal changes observed in GTA instructional practices are not the result of GTAs simply gaining more teaching experience, but are related to their experiences in the training program. These experiences include both the intentional aspects of the program (e.g. drills and feedback), as well as unintended exposure to alternate conceptions of appropriate teaching behaviors (e.g. via conversations with other GTAs and interactions with students).

Most GTAs adopted, at least temporarily, the five drilled target techniques. However, the techniques which were discussed and modeled in training sessions but not drilled were nearly completely absent from the observed classrooms (**Table 4**). These techniques (No Apology, Normalize Error, Praise Effort and Praise Improvement) are all components of the apprehension reduction dimension of active learning (Eddy, Converse, and Wenderoth 2015), which focuses on reducing students’ fear of participation and thereby lowering the barrier to a highly participatory classroom. We speculate that our failure to emphasize these techniques in training, may have hampered GTA adoption of enforced participation techniques, due to the perception (whether by GTAs or students) that enforced participation was threatening. In addition, the lack of adoption by our GTAs of techniques which were discussed and modeled, but never practiced, suggests that current methods of GTA training focusing on either literature-based or modeling-based exposure to instructional practices are not sufficient to transform GTA-led instruction.

## Conclusions and Future Directions

Our results suggest that, although GTAs are capable of utilizing evidence-based instructional practices given substantive practice-based training and regular guidance, adoption of these practices can be unstable and dependent on factors outside of the training program. This work can provide a starting model for practice-based training of graduate teaching assistants, but further work is required to understand how existing GTA attitudes towards teaching and interactions among GTAs and students influence adoption of evidence-based teaching behaviors.

Our observation that GTAs did not use non-drilled techniques suggests that discussing and modeling instructional behaviors is insufficient for modifying GTA classroom practices. Just as student mastery of academic content is aided by practice (e.g. Freeman et al. 2014), so too targeted practice appears to be a pre-requisite for mastery of an instructional skill-set. This result has implications for revising current GTA training practices, which often do not provide opportunities for practice.

For techniques which were practiced, we found that adoption varied depending on the specific instructional practice being targeted. Some techniques were adopted readily and consistently and were easily influenced by specific, goal-oriented, and timely feedback. Other practices (primarily those involved in participation enforcement) were not stably adopted and proved insensitive to formal feedback. Thus, the effectiveness of formal feedback programs for instruction may be dependent upon the particular instructional practices being targeted. We suggest future work focus on understanding the complex relationship between attitudes (both of students and of instructors) towards evidence-based teaching practices, particularly enforced participation, and instructor readiness to adopt such techniques.

## Limitations

As with all classroom-based educational studies, this work was carried out in a specific instructional setting and was likely influenced by the institutional culture present in this setting. The pre-existing structure of this course greatly facilitated our study, as GTAs for this course were already expected to attend weekly training sessions and thus our training program did not increase overall GTA time commitment. In the absence of established training requirements, introduction of a training program may cause issues with GTA buy-in. Our course also has a dedicated full-time staff member who is responsible for training and overseeing GTAs (author EJE). The existence of this resource enabled the time-intensive, repeated in-person classroom observations and one-on-one meetings called for in our training program. Courses lacking this resource may have difficulty in implementing a similar training program.

The generalizability of our study is limited by the lack of a formal control group or measurement of GTAs’ instructional practices before the beginning of the training program. Although in some institutional settings, the practices we observed may be developed organically, without training, it is the experience of our GTA coordinator (author EJE) from years of classroom observations, that GTAs for our course do not spontaneously practice these behaviors. In environments where GTAs naturally use evidence-based teaching practices, a training program such as the one described in this work would not be needed.

## Acknowledgements

The authors would like to thank the GTAs and ULAs for their patience with us in conducting this research, the undergraduate research assistants who helped transform many hours of raw video footage into data for analysis, and the many Bis2A instructors who have been instrumental in continued improvements of curricular materials for this course. We would also like to thank Dr. Marco Molinaro for his involvement in securing funding for this project.

## Supplementary Figures

**Supplementary Figure 1: Correlation matrix.** All versus all Pearson correlation matrix for technique frequency and participation levels. Correlation coefficients shown as percentages. CC = Cold Call, CCG = Group Cold Call, CCI = Individual Cold Call, Cir_Act = Circulate (active), Cir_Mod = Circulate (moderate), Cir_Pass = Circulate (passive), P = participation, RiR = Right is Right, SI.EL = Stretch it: Explain Logic, SI.FU = Stretch it: Follow-up, SQ = Student Questions, V = Volunteer, VG = Group Volunteer, VI = Individual Volunteer.

## Supplementary Tables

**Supplementary Table 1: Discussion topics.** Topics of discussion by week of term.

**Supplementary Table 2: Inter-class correlations.** Inter-class correlation values calculated for each code for the set of paired-checked (n = 23) and paired-independent (n=19) videos. Qualitative ratings for ICC values follows (Cicchetti 1994). Changes in qualitative ratings between paired-checked and paired-independent videos shown. Whether each code is included in analysis of GTA practices and/or student learning outcomes is shown.

**Supplementary Table 3: Student demographics.** Comparison of student demographic makeup for the two training datasets (Fall 2013 and Fall 2015) and the study term (Fall 2014). Final demographics of students included in model for study term also shown. First Gen. = first generation student, URM = under-represented minority.

**Supplementary Table 4: Target technique five-number summaries.** Only GTA activities (not ULA) are included in counts. Passive Circulation (in which GTA moved throughout the room, but did not interact with students) is excluded. Counts represent sums from all 158 classroom sessions in which a GTA acted as the primary instructor (excluding two observations in which a ULA acted as instructor). Five-number summaries are reported with data grouped by GTA (top) or by classroom (bottom). As each GTA led three discussion classes, the top table shows averages across those three classrooms. Min = minimum observed value, max = maximum observed value, Q1 = first quartile, Q3 = third quartile.

**Supplementary Table 5: Descriptive statistics of participation.** Five number summaries for each coded participation type reported with data grouped by GTA (top) or by classroom (bottom). As each GTA led three discussion classes, the top table shows averages across those three classrooms. Min = minimum observed value, max = maximum observed value, Q1 = first quartile, Q3 = third quartile, part = participation, Qs = questions, Indiv = individual.

**Supplementary Table 6: Longitudinal changes in classroom practices.** Differences in GTA classroom practices between first and last observations (weeks 2 and 10). Categorical magnitudes of difference correspond to threshold values provided by (Romano et al. 2006) as implemented in (Torchiano 2015) and are only shown for items where 95% confidence interval does not cross zero. These six items are shown in greater detail in **Figure 2**. Negative values of Cliff s *d* indicate decrease in frequency of practice, positive values indicate increase in frequency. For more information about interpreting Cliff s *d* values, see Methods.

**Supplementary Table 7: GTA exit survey.** Number of responses in each category for each question. Scale for section A of survey ranged from 1 = “very unhelpful” to 5 = “very helpful”, scale for sections B and C ranged from 1 = “strongly disagree” to 5 = “strongly agree”. NA option not provided for questions in section A. For all sections, 3 = “neutral”.

## Supplementary Documents

Supplementary Document 1: GTA information packet.

Supplementary Document 2: Classroom observation protocol (in-person).

Supplementary Document 3: GTA self-assessment of teaching.

Supplementary Document 4: Classroom observation protocol (video).

Supplementary Document 5: Coding manual for video observations.

Supplementary Document 6: GTA exit survey.

Supplementary Document 7: Boxplots for all technique comparisons.

